# Cell-type-specific promoters for *C. elegans* glia

**DOI:** 10.1101/2020.06.01.128546

**Authors:** Wendy Fung, Leigh Wexler, Maxwell G. Heiman

## Abstract

Glia shape the development and function of the *C. elegans* nervous system, especially its sense organs and central neuropil (nerve ring). Cell-type-specific promoters allow investigators to label or manipulate individual glial cell types, and therefore provide a key tool for deciphering glial function. In this technical resource, we compare the specificity, brightness, and consistency of cell-type-specific promoters for *C. elegans* glia. We identify a set of promoters for the study of seven glial cell types (*F16F9.3*, amphid and phasmid sheath glia; *F11C7.2*, amphid sheath glia only; *grl-2*, amphid and phasmid socket glia; *hlh-17*, cephalic (CEP) sheath glia; and *grl-18*, inner labial (IL) socket glia) as well as a pan-glial promoter (*mir-228*). We compare these promoters to promoters that are expressed more variably in combinations of glial cell types (*delm-1* and *itx-1*). We note that the expression of some promoters depends on external conditions or the internal state of the organism, such as developmental stage, suggesting glial plasticity. Finally, we demonstrate an approach for prospectively identifying cell-type-specific glial promoters using existing single-cell sequencing data, and we use this approach to identify two novel promoters specific to IL socket glia (*col-53* and *col-177*).

## INTRODUCTION

Cell-type-specific promoters are the key to studying any individual cell in *C. elegans*. With such a promoter in hand, one can visualize the cell in live animals by expressing a soluble fluorescent protein; probe cell biology in real time by expressing fluorescent constructs that label subcellular structures or serve as biosensors of cellular activity; disrupt cell function by genetic ablation or cell-specific depletion of single proteins; and determine cell-autonomy of gene function by expressing a rescuing transgene in specific cells in an otherwise mutant background.

*C. elegans* glia illustrate the power of cell-specific promoters to open up a new area of study. The powerful approaches listed above have enabled the study of several glial cell types. In particular, the amphid sheath (AMsh) and cephalic sheath (CEPsh) glia have been studied in detail over the last two decades, while more recent work has begun to examine the amphid socket (AMso), phasmid socket (PHso1 and PHso2), and inner labial socket (ILso) glia (Singhvi and Shaham, 2019). In contrast, other glial cell types, for which no cell-type-specific promoters have been identified, remain largely unexplored.

In this technical resource, we summarize and compare the promoters that have been used to study distinct glial cell types in *C. elegans*. We evaluate a set of specific, bright, and consistent promoters for the study of seven glial cell types as well as a pan-glial driver. We note that some promoters exhibit interesting dynamics in their expression patterns that are suggestive of glial plasticity. Finally, we demonstrate an approach to identify new cell-type-specific glial promoters by taking advantage of recent single-cell sequencing experiments, and we use this approach to identify *col-53* and *col-177* as novel cell-specific promoters for ILso glia.

## RESULTS AND DISCUSSION

### Survey of glial promoters used in the literature

We performed a survey of cell-specific promoters that have been used to label and manipulate *C. elegans* glia (Table 1). All *C. elegans* glia are associated with sense organs, mostly in the head and tail (Ward et al., 1975). Hermaphrodites have 24 sense organs: 18 in the head (two amphid, AM; four cephalic, CEP; six inner labial, IL; six outer labial, OL), two in the anterior midbody (anterior deirid, ADE); two in the posterior midbody (posterior deirid, PDE), and two in the tail (phasmid, PH) (White et al., 1986). Each sense organ comprises one or more ciliated sensory neurons; one sheath glial cell; and, in most cases, one socket glial cell (the phasmid has two socket glia). The distal endings of the sheath and socket glia are arranged to form a tube-shaped epithelium that is continuous with the skin (Fig. 1) (Low et al., 2019; Ward et al., 1975). The proximal portion of this tube is formed by the ending of the sheath glial cell, while the distal portion of the tube is formed by the ending of the socket glial cell. In each sense organ, sensory neurons protrude through this glial tube to directly access the environment (Fig. 1). The sheath (sh) and socket (so) glia of most sense organs appear to represent distinct cell types. For example, the AMsh glia are a different cell type than the AMso glia or CEPsh glia based on morphology, molecular profile, and function (Mizeracka and Heiman, 2015). As discussed in detail below, each of these glial cell types expresses a distinct transcriptional reporter. The functional relevance of these striking transcriptional differences has not been fully established, but these transcriptional reporters serve as powerful tools to study and manipulate individual glial cell types. In addition to the glial cell types described above, there are also sex-specific glia in copulatory sense organs in the male tail, and glial-like functions can be performed by mesoderm-derived GLR cells and by the skin (Singhvi and Shaham, 2019; Yang and Chien, 2019). Here, we will focus exclusively on sheath and socket glia of sense organs that are present in both sexes.

**Table 1.** Glial promoters used in the literature. Promoters with glial-specific expression were identified in the literature. Within each group, promoters are listed in order from most to least frequently used.

| **AMsh or PHsh** |  |  |
| --- | --- | --- |
| *F16F9.3* | 2 kb promoter | (Bacaj et al., 2008; Braunreiter et al., 2014; Heiman and Shaham, 2009; Low et al., 2019; Mizeracka et al., 2019; Oikonomou et al., 2011; Procko et al., 2012, 2011; Singhvi et al., 2016; Wallace et al., 2016; Yip and Heiman, 2018) |
| *vap-1*<br>(developmentally regulated) | 2797 bp or 5205 bp promoter | (Bacaj et al., 2008; Ding et al., 2015; Oikonomou et al., 2011; Perens and Shaham, 2005; Procko et al., 2011; Wallace et al., 2016; Wang et al., 2017, 2012, 2008) |
| *T02B11.3* | 2.5 kb promoter | (Bacaj et al., 2008; Grant et al., 2015; Mizeracka et al., 2019; Oikonomou et al., 2012, 2011; Procko et al., 2011; Wang et al., 2017, 2008) |
| *fig-1* | 2.2 kb promoter | (Bacaj et al., 2008; Fenk and de Bono, 2015; Kage-Nakadai et al., 2016; Wallace et al., 2016; Yoshida et al., 2016) |
| *ver-1*<br>(thermo- and dauer-regulated) | 2110 bp promoter | (Popovici et al., 2002; Procko et al., 2012, 2011; Yoshida et al., 2016) |
| *F53F4.13* | 650 bp promoter | (Bacaj et al., 2008; Mizeracka et al., 2019; Singhvi et al., 2016) |
| *F11C7.2* | 350 bp promoter | (Bacaj et al., 2008; Wallace et al., 2016) |
| *hmit-1.2*<br>(osmo-regulated) | 979 bp promoter | (Kage-Nakadai et al., 2016, 2011) |
| *lit-1* | 2.5 kb promoter | (Oikonomou et al., 2011; Wallace et al., 2016) |
| *grl-28* | 2553 bp promoter | (Hao et al., 2006) |
| *acd-1* | 2550 bp promoter, plus coding sequences | (Wang et al., 2008) |
| *F52E1.2* | 3.3 kb promoter | (Ohkura and Bürglin, 2011) |
| *hmit-1.3* | 4883 bp promoter | (Kage-Nakadai et al., 2011) |
| *F58F9.6* | 4 kb promoter | (Procko et al., 2012) |
| *K02E11.4* | 800 bp promoter | (Wallace et al., 2016) |
| *R11D1.3* | 900 bp promoter | (Wallace et al., 2016) |
| *bgnt-1.1* | recombineered fosmid | (Timbers et al., 2016) |
| **AMso or PHso** |  |  |
| *itr-1* | 2.3 kb promoter (from between exon 1 and exon 2) | (Gower et al., 2001; Han et al., 2013; Heiman and Shaham, 2009; Mizeracka et al., 2019; Sammut et al., 2015; Tucker et al., 2005; Wang et al., 2017) |
| *grl-2* | 2981 bp promoter | (Hao et al., 2006; Hunt-Newbury et al., 2007; Low et al., 2019; Mizeracka et al., 2019; Molina-Garcia et al., 2018; Sammut et al., 2015) |
| *lin-48* | 6.8 kb promoter | (Johnson et al., 2001; Mizeracka et al., 2019; Molina-Garcia et al., 2018) |
| *grd-15* | 840 bp promoter | (Hunt-Newbury et al., 2007; Timbers et al., 2016) |
| *unc-53* | 3.4 kb promoter (from exon 8 to exon 13) | (Stringham et al., 2002; Tucker et al., 2005) |
| *alr-1* | 1 kb promoter | (Tucker et al., 2005) |
| *abt-4* | 2472 bp promoter | (Hunt-Newbury et al., 2007) |
| *grl-6* | 939 bp promoter | (Hunt-Newbury et al., 2007) |
| *srp-2* | 2833 bp promoter | (Hunt-Newbury et al., 2007) |
| **Combinations of AMsh, AMso, PHsh, and PHso** |  |  |
| *daf-6* | 3 kb promoter | (Kage-Nakadai et al., 2016; Moussaif and Sze, 2009; Ohkura and Bürglin, 2011; Perens and Shaham, 2005; Wallace et al., 2016) |
| *ztf-16* | 2.1 kb enhancer (from 4637 to 2536 bp upstream of ATG) | (Procko et al., 2012; Sammut et al., 2015) |
| *ttx-1* | 3.5 kb enhancer (from 11 to 7.5 kb upstream of ATG) | (Procko et al., 2011) |
| **CEPsh** |  |  |
| *hlh-17* | 2.5 kb promoter or 1.9 kb promoter | (Cianciulli et al., 2019; Colón-Ramos et al., 2007; Gibson et al., 2018; Ji et al., 2019; Katz et al., 2018; McMiller and Johnson, 2005; Mizeracka et al., 2019; Rapti et al., 2017; Sammut et al., 2015; Shao et al., 2013; Stout and Parpura, 2011; Yoshida et al., 2016; Yoshimura et al., 2008) |
| *swip-10* | 1.5 kb enhancer (from fifth intron, 1013 to 2504 bp downstream of ATG) | (Hardaway et al., 2015) |
| *twk-16* | 3412 bp promoter | (Cianciulli et al., 2019) |
| **ILso** |  |  |
| *grl-18* | 2968 or 2962 bp promoter | (Cebul et al., 2020; Hao et al., 2006; Mizeracka et al., 2019) |
| **Combinations of ILsh, ILso, OLsh, and OLso** |  |  |
| *itx-1* | 2977 bp promoter or 1.8 kb promoter | (Haklai-Topper et al., 2011; Han et al., 2013; McQuary et al., 2016; Sammut et al., 2015) |
| *delm-1* | 2 kb promoter | (Han et al., 2013) |
| *delm-2* | 1.4 kb promoter | (Han et al., 2013) |
| **Most (or many) glia** |  |  |
| *ptr-10* | 300 bp promoter | (Gibson et al., 2018; Hardaway et al., 2015; Katz et al., 2018; Oikonomou et al., 2011; Rapti et al., 2017; Sammut et al., 2015; Wallace et al., 2016; Yin et al., 2017; Yoshida et al., 2016; Yoshimura et al., 2008) |
| *mir-228* | 2.2 kb promoter | (Molina-García et al., 2018; Pierce et al., 2008; Rapti et al., 2017; Wallace et al., 2016; Wang et al., 2017; Yin et al., 2017) |
| *kcc-3* | 3.9 kb promoter and 2.2 kb downstream sequences | (McQuary et al., 2016; Singhvi et al., 2016; Tanis et al., 2009; Yoshida et al., 2016) |
| *pros-1* | recombineered fosmid or 9 kb promoter | (Kage-Nakadai et al., 2016; Wallace et al., 2016) |
| **Unidentified combinations of glia** |  |  |
| *grd-8* | 905 bp promoter | (Hao et al., 2006) |
| *grd-16* | 2866 bp promoter | (Hao et al., 2006) |
| *grl-1* | 287 bp promoter | (Hao et al., 2006) |
| *grl-6* | 940 bp promoter | (Hao et al., 2006) |
| *grl-8* | 2893 bp promoter | (Hao et al., 2006) |
| *grl-12* | 2989 bp promoter | (Hao et al., 2006) |
| *grl-13* | 2843 bp promoter | (Hao et al., 2006) |
| *grl-17* | 2869 bp promoter | (Hao et al., 2006) |
| *grl-18* | 2905 bp promoter | (Hao et al., 2006) |
| *grl-29* | 1893 bp promoter | (Hao et al., 2006) |
| *rgba-1* | 2.1 kb promoter | (Yin et al., 2017) |
| *argk-1* | 1.2 kb promoter | (McQuary et al., 2016) |

**Fig. 1.**
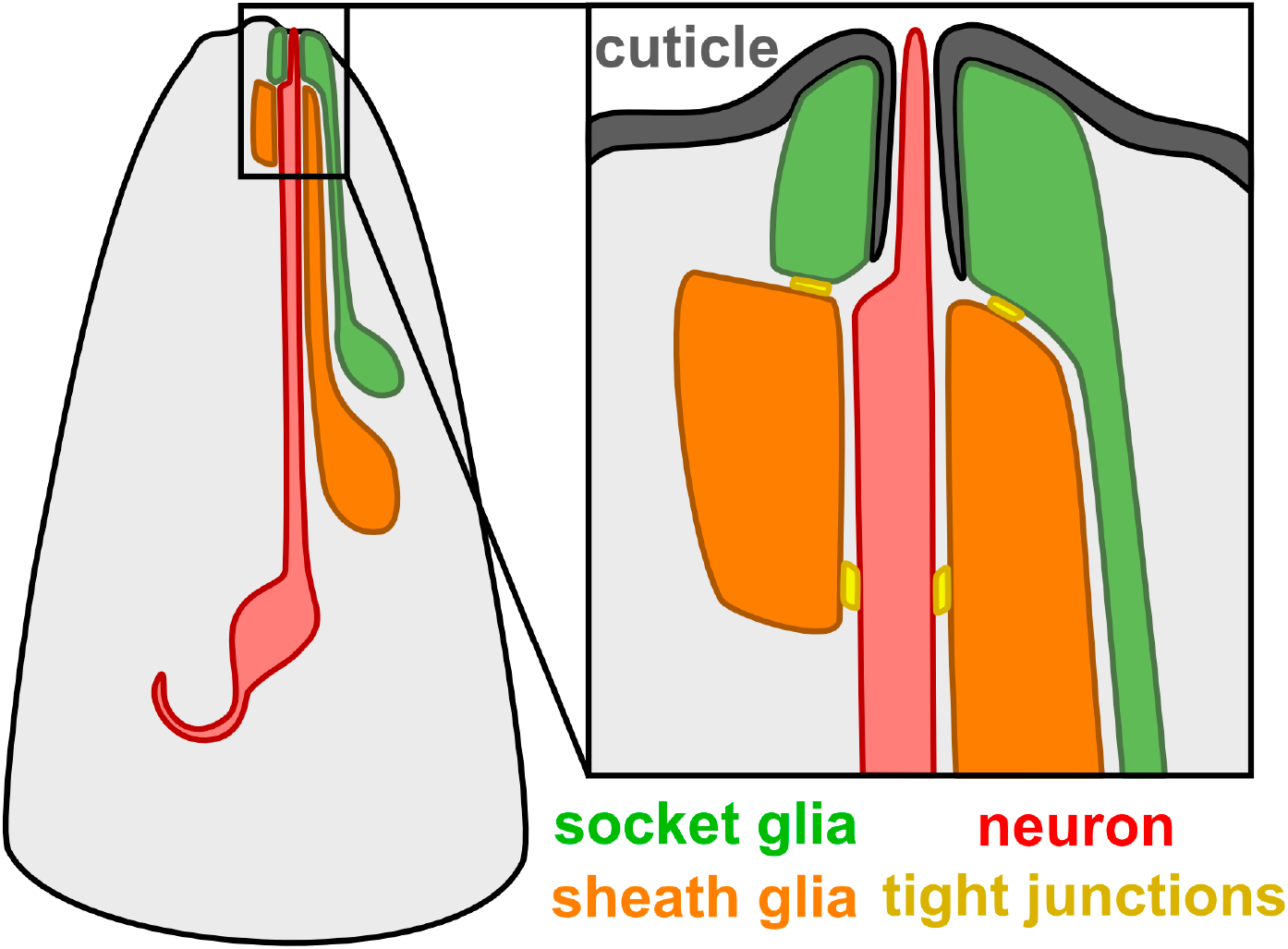
Schematic of a *C. elegans* sense organ in the head. A typical *C. elegans* sense organ contains one or more sensory neurons (red) and two glial cells, called the sheath (orange) and socket (green). Most sensory neurons extend a ciliated dendritic ending through a tube-shaped pore formed by the sheath and socket glia. The socket glia secretes cuticle (gray) that forms an open pore through which chemosensory dendrite endings protrude, as shown, or a closed sheet into which mechanosensory dendrite endings are embedded (not shown). Tight junctions (yellow) are present between the neuron and sheath glia, the sheath and socket glia, and the socket glia and skin.

Cell-type-specific promoters for *C. elegans* glia have been identified through three approaches: reverse genetics, forward genetics, and transcriptional profiling. In reverse genetics, the molecular identity of a gene motivates investigation of its expression pattern. The study of molecularly interesting gene families led to the first identification of promoters expressed specifically in glia, including the basic helix-loop-helix transcription factor gene *hlh-17* (McMiller and Johnson, 2005; Yoshimura et al., 2008); the inositol triphosphate receptor gene *itr-1* (Gower et al., 2001); the neurexin gene *itx-1* (Haklai-Topper et al., 2011); the VEGF receptor gene *ver-1* (Popovici et al., 2002); the venom allergen protein gene *vap-1* (Bacaj et al., 2008); the ion channel genes *delm-1* and *delm-2* (Han et al., 2013); the microRNA *mir-228* (Pierce et al., 2008); and a group of nematode-specific genes with distant similarity to Hedgehog genes, including the "groundhog" (*grd*) and "ground-like" (*grl*) genes (Hao et al., 2006). In a typical experiment, a putative promoter fragment from the gene of interest is inserted upstream of GFP in a transgene and the resulting expression pattern is examined. Expression in the AMsh (*vap-1*, *ver-1*) or CEPsh (*hlh-17*) glia is readily recognized due to the symmetry and distinctive morphology of these cells (Table 1). However, expression in the ILsh, ILso, OLsh, and OLso glia is harder to recognize due to the variable cell body positions and similar symmetry and morphology of these glial types (Bargmann and Avery, 1995). Thus, most promoters that label combinations of these cells, including *itx-1* and the *delm, grl*, and *grd* gene families, have not had their expression patterns precisely defined (Table 1).

As a second approach, forward genetics has led to the discovery of glial-expressed genes through the identification of mutants that disrupt neuronal or glial function. These include *daf-6* and *lit-1*, which affect glial morphogenesis and thus sensory neuron function (Oikonomou et al., 2011; Perens and Shaham, 2005); *ttx-1* and *ztf-16*, which affect a glial-specific temperature response (Procko et al., 2012, 2011); and *kcc-3*, which affects the microenvironment that glia create around sensory neuron endings (Singhvi et al., 2016; Yoshida et al., 2016) (Table 1). These genes came under study due to their roles in AMsh glia, but several are also expressed in other classes of glia.

Finally, using transcriptional profiling approaches, existing glial markers have been used to “bootstrap” the way to new markers, typically by using fluorescence-activated cell sorting to purify a glial population and then subjecting it to RNA profiling. This approach led to the identification of numerous AMsh markers, including *fig-1*, *F16F9.3*, *T02B11.3*, and others (Bacaj et al., 2008; Wallace et al., 2016) (Table 1). The expression of these genes is highly specific to the AMsh, but they were missed by other approaches because they are not members of a conserved gene family and (with the exception of *fig-1*) have not been found to cause phenotypes when disrupted. The AMsh and PHsh are highly similar, and many AMsh markers are also expressed in the PHsh.

### Recommended promoters for specific glial cell types

There is currently no consensus on which promoters are most appropriate for studying specific glia types. Therefore, we collected glial promoters from the literature and reexamined the specificity, brightness, and consistency of their expression patterns. Specificity requires that expression is restricted to individual glial cell types for easy visualization and for reducing off-target effects of genetic manipulations. Brightness allows fine structural details of glia to be resolved, which provides a tool for examining glia-neuron interactions and an efficient way to screen for mutants with glial morphology defects. Last, consistency assesses whether the promoter reliably labels the same number of cells across individuals. Based on these criteria, we selected five promoters to target seven glial cell types: the AMsh and PHsh; CEPsh; the AMso, PHso1 and PHso2; and ILso glia (Fig. 2, Table 2). However, it is important to note that brightness and consistency are also affected by the structure of the transgene, including copy number and whether it is genomically integrated, as well as the fluorescent reporter used.

**Fig. 2.**
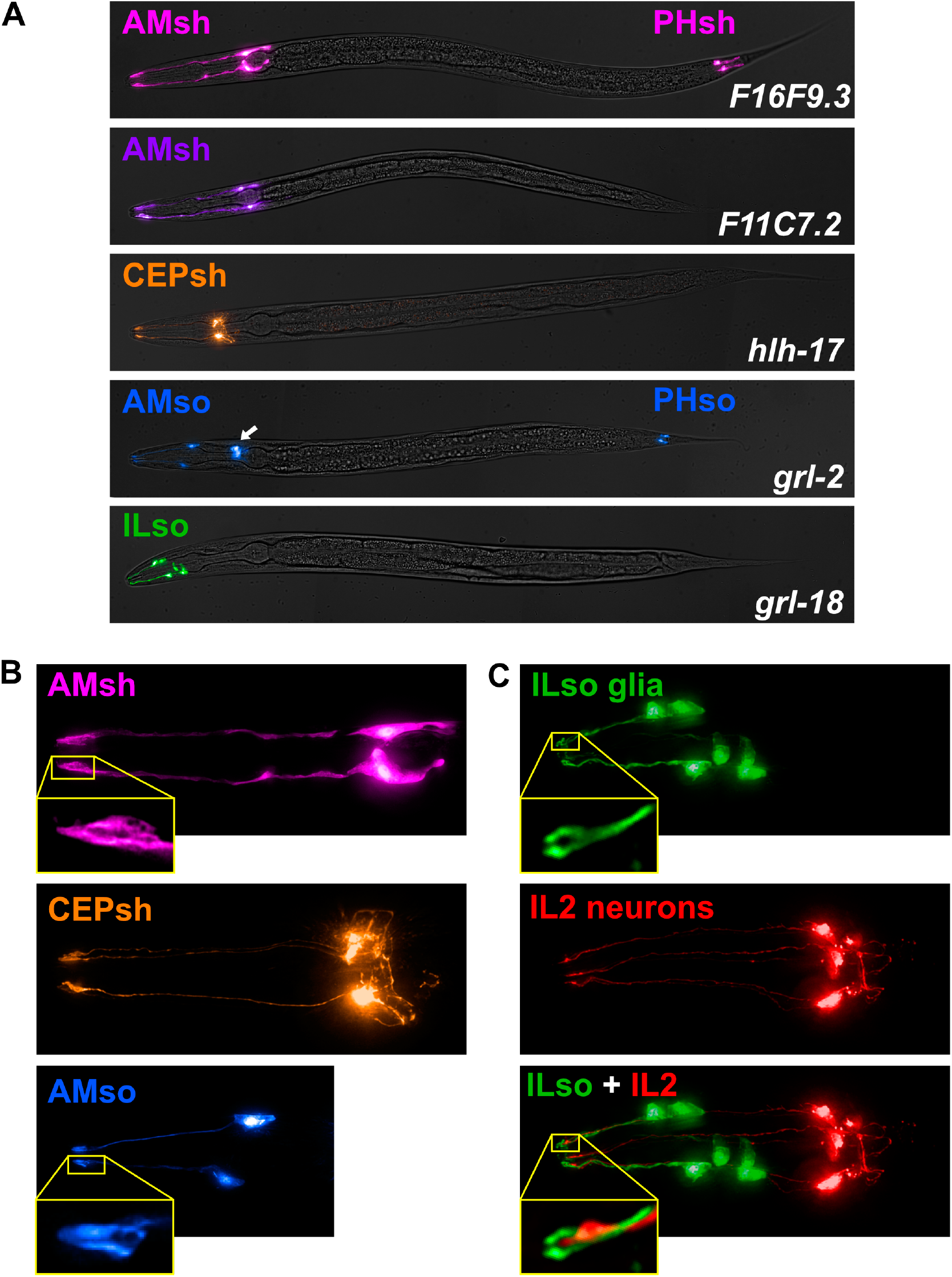
Cell-type-specific promoters for sheath and socket glia. (A) Merged brightfield images and pseudo-colored fluorescence projections of animals expressing fluorescent proteins under control of promoters selected for brightness, specificity, and consistency. *F16F9.3* (pink, AMsh and PHsh); *F11C7.2* (purple, AMsh only); *hlh-17* (orange, CEPsh); *grl-2* (blue, AMso, PHso1, and PHso2; also expressed in excretory duct and pore cells, white arrow); *grl-18* (green, ILso). (B) Magnified images of AMsh (pink, *F16F9.3*), CEPsh (orange, *hlh-17)*, and AMso (blue, *grl-2)*, showing that their brightness is sufficient to resolve fine structural details including the tube-like pores of the AMsh and AMso glia and the branch-like posterior processes of the CEPsh glia. (C) Head of an animal expressing *grl-18* pro:GFP (green, ILso glia) and *klp-6* pro:mCherry (red, IL2 neurons), demonstrating that the *grl-18* promoter labels the six ILso glia, identified by their processes which each form a pore for the ciliated dendritic ending of an IL2 sensory neuron.

**Table 2.** Recommended cell-type-specific glial promoters. For each marker, the promoter column indicates the size (bp) of the genomic DNA fragment that is taken directly upstream of the gene translation start site, except *hlh-17* for which the 3’ end of the promoter fragment is 118 bp upstream of the translation start site.

| Gene | Promoter (bp) | Glia marked | Other cells marked | References |
| --- | --- | --- | --- | --- |
| *F16F9.3* | 2000 | AMsh, PHsh | none | (Bacaj et al., 2008) |
| *F11C7.2* | 350 | AMsh | none | (Bacaj et al., 2008) |
| *hlh-17* | 2500 | CEPsh | none | (McMiller and Johnson, 2005;<br>Yoshimura et al., 2008) |
| *grl-2* | 2981 | AMso, PHso1, PHso2 | excretory duct, pore | (Hao et al., 2006) |
| *grl-18* | 2962 | ILso | vulval epithelial cells | (Cebul et al., 2020; Hao et al., 2006) |
| *mir-228* | 2200 | all (or nearly all) glia | seam cells, excretory cell | (Pierce et al., 2008) |

A pair of AMsh glia are part of the bilateral amphid sense organs in the head (Ward et al., 1975). Each AMsh is associated with 12 sensory neurons that respond to diverse stimuli including chemical and mechanical cues, temperature, osmolarity, and pheromones (Bargmann, 2006). AMsh glia are the most well-studied *C. elegans* glia, with many highly specific markers identified (Bacaj et al., 2008; Wallace et al., 2016) (Table 1). We selected *F16F9.3* and *F11C7.2* as AMsh promoters that offer different advantages (Fig. 2A). *F16F9.3* is reliably expressed in AMsh glia as well as the PHsh glia, their functional counterparts in the tail (24/25 animals with an integrated *F16F9.3*pro:mCherry transgene had 2/2 AMsh and 2/2 PHsh glia marked; 1/25 animals had 1/2 AMsh and 2/2 PHsh glia marked) (Fig. 2A, Supp. Table S2). The strong expression of *F16F9.3* allows us to clearly visualize the tube-shaped pores in the sheath glia through which the sensory dendrite endings of the neurons protrude (Fig. 2B). Other promoters, including *F53F4.13* and *T02B11.3*, also offer strong specific labeling of AMsh and PHsh. By comparison, because *F11C7.2* expression is restricted to the AMsh glia (Fig. 2A), it offers the advantage of genetically manipulating AMsh glia without affecting PHsh glia (16/23 animals with an *F11C7.2*pro:GFP extrachromosomal array had 2/2 AMsh glia and no PHsh glia marked, whereas 7/23 animals had 1/2 AMsh glia and no PHsh glia marked, Supp. Table S2). In summary, *F16F9.3* has higher overall expression, while *F11C7.2* has higher specificity for AMsh glia.

Each of four CEPsh glia is associated with the four-fold symmetric cephalic sense organs in the head. Each CEPsh glial cell wraps around the dendrite of a mechanosensory neuron (CEP) and, in males, an additional pheromone-detecting neuron (CEM) (Ward et al., 1975). We selected *hlh-17* as a cell-specific promoter for CEPsh glia (Fig. 2A). We evaluated the expression pattern of a transcriptional reporter for *hlh-17* (McMiller and Johnson, 2005; Yoshimura et al., 2008) and observed that it is restricted to CEPsh glia and is consistently expressed in all four cells (22/22 animals with an integrated *hlh-17*pro:GFP transgene had 4/4 CEPsh glia marked, Supp. Table S2). Though the transcriptional reporter we examined uses a ~2.5 kb promoter fragment and appears to be highly specific to CEPsh glia, it is important to note that reporters containing a different promoter fragment of the *hlh-17* gene are expressed in additional cells (Yoshimura et al., 2008). The marker typically becomes brighter as animals reach adulthood but the extended anterior and posterior processes of the glia are visible at all larval stages. We can clearly observe the thin, branch-like posterior processes that wrap around the nerve ring – a feature that distinguishes CEPsh glia from other glia of *C. elegans* (Fig. 2B). The ability to visualize these fine details makes it possible to investigate the role of CEPsh glia in shaping neuronal connections in the central neuropil, or nerve ring (Colón-Ramos et al., 2007; Ji et al., 2019; Rapti et al., 2017; Shao et al., 2013).

The two AMso glia wrap around the distal tips of a subset of amphid sensory neurons and guide them towards pores in the overlying cuticle so that they can access the external environment (Ward et al., 1975). We selected *grl-2* to label and target AMso glia (Fig. 2A). It is consistently expressed in AMso glia as well as the PHso glia in the tail (23/23 animals with an integrated *grl-2*pro:YFP transgene have 2/2 AMso glia, 2/2 PHso1, and 2/2 PHso2 glia marked, Supp. Table S2). However, the marker is also expressed brightly in the excretory duct and pore cells (Hao et al., 2006) (Fig. 2A, arrow), which makes it problematic to use *grl-2* for genetic perturbations, as disrupting the excretory cells can lead to lethality or defects in osmoregulation (Nelson and Riddle, 1984; Sundaram and Buechner, 2016). Although *grl-2* is less cell-type-specific than the AMsh or CEPsh promoters, it is bright (Fig. 2B) and consistent and exhibits a more restricted expression pattern than other AMso glia markers that have been identified to date (Table 1). As previous work has shown, we found that *lin-48* offers more restricted expression in the phasmid glia (PHso1 but not PHso2) but is also expressed in the posterior intestine and additional cells in the head, including the excretory duct and unidentified neurons (Johnson et al., 2001). It will be important to identify new AMso glia markers whose expression is excluded from the excretory cells so that they can be used as tools to elucidate AMso glia biology and function in future studies.

Last, we selected the *grl-18* promoter for the six ILso glia that are part of the six-fold symmetric inner labial sense organs of the head (Fig. 2A). In a previous study (Hao et al., 2006), *grl-18* was tentatively identified as a potential marker for ILso or the OLso quadrant (dorsal and ventral, but not lateral) glia based on position and morphology. We find that *grl-18* is a highly specific marker for ILso glia. Each ILso glia wraps around the dendrite endings of two sensory neurons, called the IL1 and IL2 neurons (Ward et al., 1975). Therefore, we examined animals co-expressing *grl-18*pro:GFP and the IL2 neuron-specific marker *klp-6*pro:mCherry (Fig. 2C). We observed the ciliated endings of the IL2 dendrites protruding through the tube-like pores formed by the glia expressing *grl-18*pro:GFP, demonstrating that *grl-18* labels ILso glia. This is consistent with our previous observation that *grl-18*pro:GFP labels the lateral ILso glial cells that form specialized attachments to the BAG and URX neurons (Cebul et al., 2020). To assess the quality of this marker, we examined the expression pattern more carefully. We found that its expression is highly consistent (28/28 animals with an integrated *grl-18*pro:GFP transgene had 6/6 ILso glia marked, Supp. Table S2). It is mostly specific to the ILso glia, however after animals enter the fourth larval (L4) stage, *grl-18*pro:GFP is also expressed in vulval epithelial cells. The marker is very bright, so it can be used to visualize the pore structure of the glia (Fig. 2C) and screen for mutants with defects in morphology to investigate ILso glia development. It is also suitable for structured illumination microscopy, which requires greater brightness than conventional imaging (Cebul et al., 2020). An inherent limitation is that the ILso cell body positions are variable (Bargmann and Avery, 1995) and often appear to overlap in a two-dimensional projection, making it difficult to show all six cells clearly in a single image. Overall, we have demonstrated that *grl-18* specifically labels ILso glia, and this opens up opportunities for studying ILso glia, which remain largely unexplored.

Based on their specificity, brightness, and consistency, the markers listed in Table 2 are especially useful for labeling and genetically manipulating the AMsh, PHsh, CEPsh, AMso, PHso1, PHso2, and ILso glia. We are not aware of promoters that specifically label the CEPso, OLsh, OLso, ADEsh, ADEso, PDEsh, or PDEso. In fact, little is known about these glia at all. Currently, the best approach for studying them may be to use promoters that are expressed in combinations of multiple glial cell types, as described further below.

### Promoters that label multiple glial cell types

Pan-neuronal promoters, such as *rab-3*pro (Nonet et al., 1997), have proven vital for the study of neuronal function and, similarly, the ability to drive expression in all glial cells is important for studying glial function. Several promoters that drive expression in many glial types have been identified (Table 1). We selected *mir-228* as a pan-glial promoter, because it drives robust expression in most if not all glia (Fig. 3A, C) (Pierce et al., 2008). However, it is important to note that it is also expressed in some non-glial cells, including the hypodermal seam cells and the excretory cell.

**Fig. 3.**
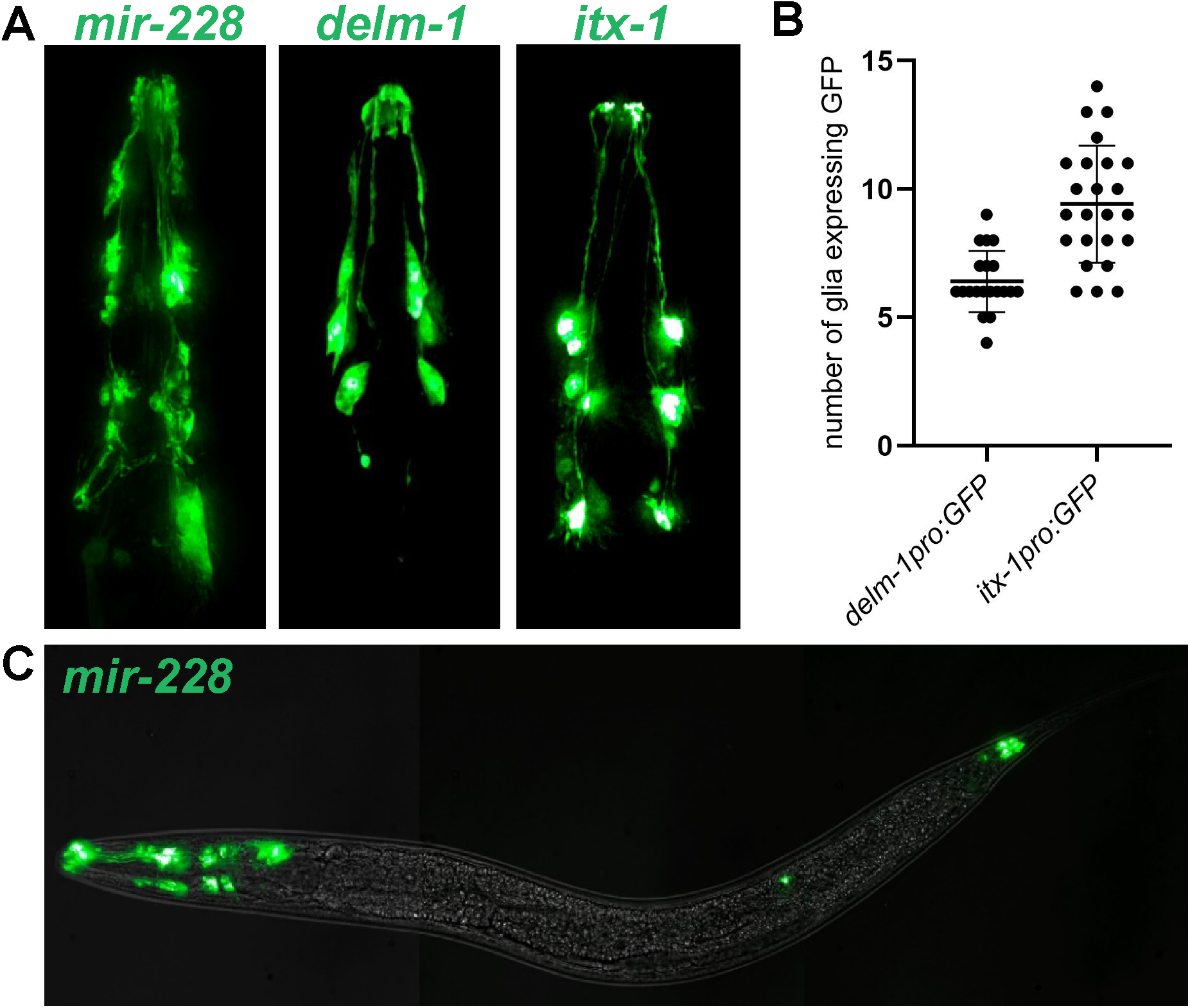
Promoters expressed in combinations of glia. (A) Images of head expression of GFP under control of the pan-glial promoter *mir-228* and the variable glial promoters *delm-1* and *itx-1*. (B) Quantification of the number of GFP-expressing glia observed with *delm-1* (n=20) and *itx-1* (n=24) extrachromosomal reporters. Error bars indicate standard deviation. *p*<0.0001, unpaired t-test with Welch’s correction. (C) Merged brightfield and fluorescence projections of a whole animal expressing pan-glial *mir-228*pro:GFP.

Promoters expressed in other combinations of glia are also useful for studying glial biology. Curiously, some of these promoters exhibit more variable expression across individuals than the highly consistent cell-type-specific promoters described above. Two such promoters are *delm-1* and *itx-1*, both of which are expressed in combinations of glia in the head (Fig. 3A) (Haklai-Topper et al., 2011; Han et al., 2013). While the identity of some of these glia is known (see Table 1), the expression patterns of these promoters have not been fully characterized. Although there is some overlap in their expression, as described previously (Han et al., 2013), it is clear that *delm-1* and *itx-1* are expressed in different combinations of glia (Fig. 3A). In contrast to the highly consistent cell-type-specific promoters described above, *delm-1*pro:GFP and *itx-1*pro:GFP extrachromosomal arrays exhibit greater variability from animal to animal, although each promoter shows a consistent range of expression (Fig. 3B). *delm-1* is expressed in fewer glia and has less variability in expression than *itx-1* (Fig. 3B, Supp. Table S2). A high degree of variability in GFP intensity between different glia within the same animal was often observed, particularly for *itx-1*. It is possible that differences in the threshold of detection introduced more variability into the number of GFP-expressing glia that could be discerned. Despite this variability, these promoters provide a useful tool for driving expression in combinations of glia that cannot be accessed with other promoters.

### Plasticity in expression of glial promoters

The variability in expression of these promoters may reflect transgene artifacts or true biological differences across individuals. Indeed, the expression of some glial promoters has been shown to be dynamic across life stages, suggesting that glia exhibit surprising plasticity in response to environmental conditions and the internal state of the animal. We noted that the expression of *vap-1* in the AMsh is dependent upon developmental stage. Using a 2797 bp promoter, an integrated *vap-1*pro:RFP transgene exhibits absent or very faint expression in the AMsh prior to the L4 stage, but it then becomes robust in L4 animals and is upregulated further in adulthood (Fig. 4A,C). These developmental changes in expression suggest that changes in glial function may occur during late larval development and maturation. In addition to developmental stage, *vap-1* expression is subject to further modulation through activity of amphid sensory neurons. DYF-7 is required to anchor the dendrite endings of sensory neurons to the tip of the nose, and loss of function of this protein results in shortened dendrites that diminish the function of these sensory neurons (Heiman and Shaham, 2009; Low et al., 2019). Loss of DYF-7 function due to the null allele *dyf-7(ns119)* causes severe defects in *vap-1* expression in AMsh glia, suggesting that input from the external environment may modulate AMsh gene expression (Fig. 4C). Interestingly, while *vap-1*pro:RFP expression at the L4 stage is nearly lost in *dyf-7(ns119)* mutants, it is still upregulated to some degree in adults, albeit to a much lesser extent than in wild type (Fig. 4C). These observations suggest that there may be multiple layers of plasticity in glial gene expression, both through developmental stage and sensory activity.

**Fig. 4.**
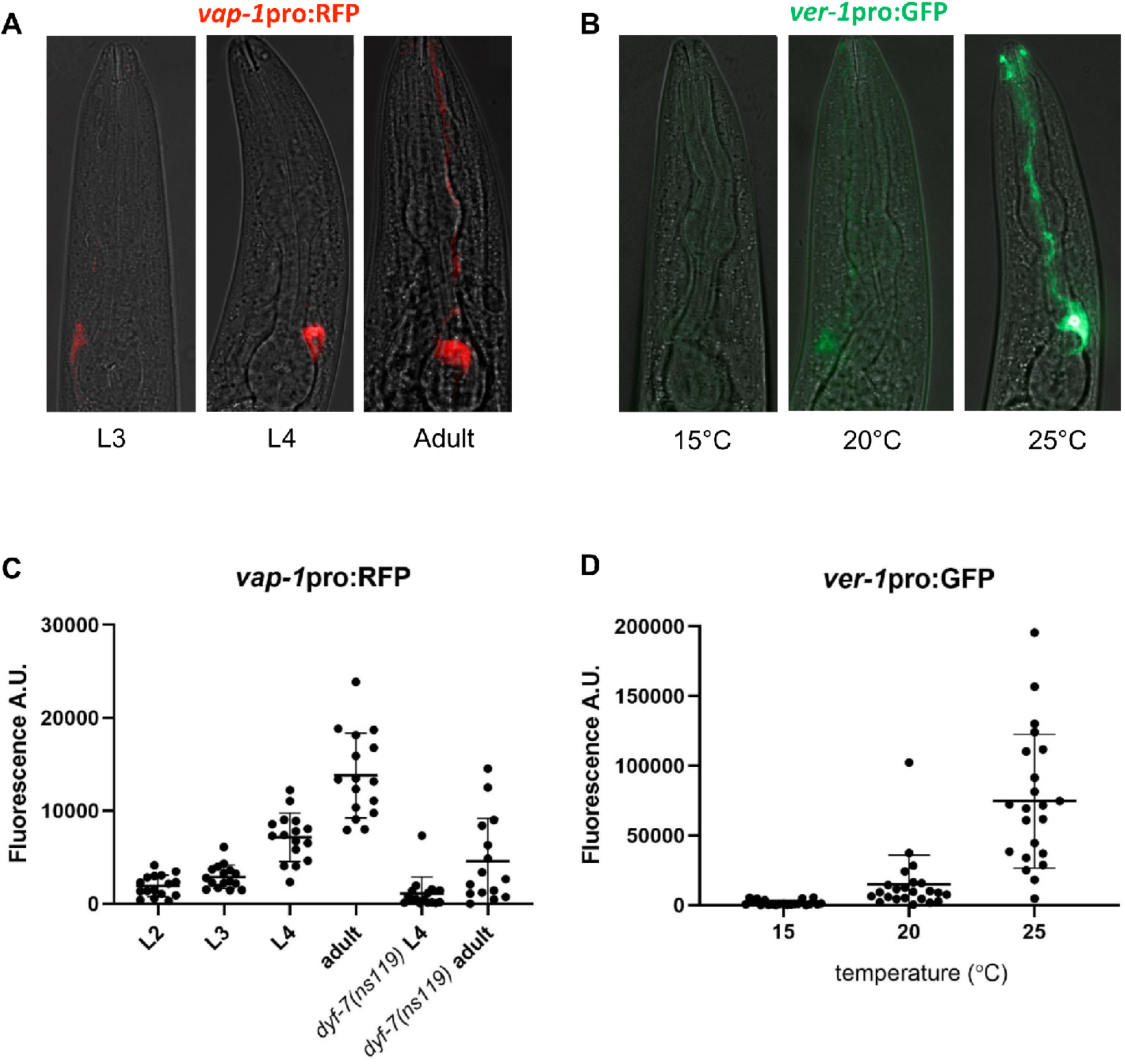
Plasticity in glial gene expression. (A) *vap-1*pro:RFP expression in AMsh glia in animals at the third larval (L3), fourth larval (L4), and 1-day adult life stages, showing developmental regulation of gene expression. (B) *ver-1*pro:GFP expression in AMsh glia in L4 animals grown at 15°C, 20°C, or 25°C, showing thermoregulation of gene expression as previously described (Procko et al., 2012, 2011). (C) Quantification of *vap-1*pro:RFP fluorescence in AMsh glia from second larval (L2) stage to 1-day adults and in *dyf-7(ns119)* L4 and adult animals, showing that sensory defects alter gene expression. n=15-16 per group. *p*<0.05, *dyf-7* L4 vs *dyf-7* adults; *p*<0.0001, *dyf-7* adults vs wild-type adults; unpaired t-test with Welch’s correction (D) Quantification of *ver-1*pro:GFP fluorescence in the AMsh glia in L4 animals from 15°C to 25°C. n=21-23 per group. *p*<0.05, 20°C vs 15°C; *p*<0.0001, 20°C vs 25°C; Brown-Forsyth and Welch one-way ANOVA with Dunnett’s multiple comparisons test.

The expression of *ver-1* in AMsh glia has been previously reported to be modulated by temperature, increasing steadily from undetectable expression at 15°C to very strong expression at 25°C (Procko et al., 2012, 2011). We were able to recapitulate these results (Fig. 4B,D). Additionally, *ver-1* expression has also been shown to be induced by entry into dauer, a long-lived stress-resistant life stage, concomitant with remodeling of glia (Procko et al., 2011).

Other examples of dynamic gene expression in glia include *hmit-1.2*, which is upregulated in AMsh glia in response to osmotic stress (Kage-Nakadai et al., 2011), and *daf-6*, which is downregulated in AMsh glia during dauer (Moussaif and Sze, 2009). While these data provide evidence for glial plasticity, the functional significance of these changes in gene expression for AMsh function and its interactions with neurons remain unclear. While all of these examples involve AMsh glia, it is likely that other glia may exhibit similar plasticity. Dynamic expression of promoters in other glia has yet to be identified and the full extent of glial plasticity in response to different environmental and state changes remains unknown.

### Single-cell transcriptional profiling identifies new cell-type-specific promoters for glia

Discovery of glial cell-type-specific promoters has often relied on serendipity, and has been limited to study of one glial cell type at a time. The recent revolution in single-cell transcriptional profiling raises the possibility that promoters specific to each glial cell type can be identified systematically. Single-cell transcriptional profiling of embryos identified progenitor cells that give rise to eight major groups of tissues, including a group that contains glia and excretory cells. Packer et al. used existing markers from the literature to propose identities for clusters of cells corresponding to nine glial subtypes (Packer et al., 2019).

In order to test whether this dataset could be used to prospectively identify novel cell-type-specific glial markers, we attempted to identify a novel marker for ILso glia. The ILso glia cluster is characterized by its enrichment in *grl-18* expression (Fig. 5A). Using the VisCello data explorer (http://github.com/qinzhu/VisCello.celegans), we generated a list of additional genes that are highly enriched in the ILso glia cluster compared to the global dataset (see Methods). From the list, we selected the collagen genes, *col-53* and *col-177*, to characterize further. They are strong candidates to be markers for ILso glia, because they are highly expressed in the ILso glia cluster (Fig. 5A, Supp. Fig. S1A), like *grl-18*, and because socket glia secrete cuticle, which consists of collagen and other extracellular matrix proteins.

**Fig. 5.**
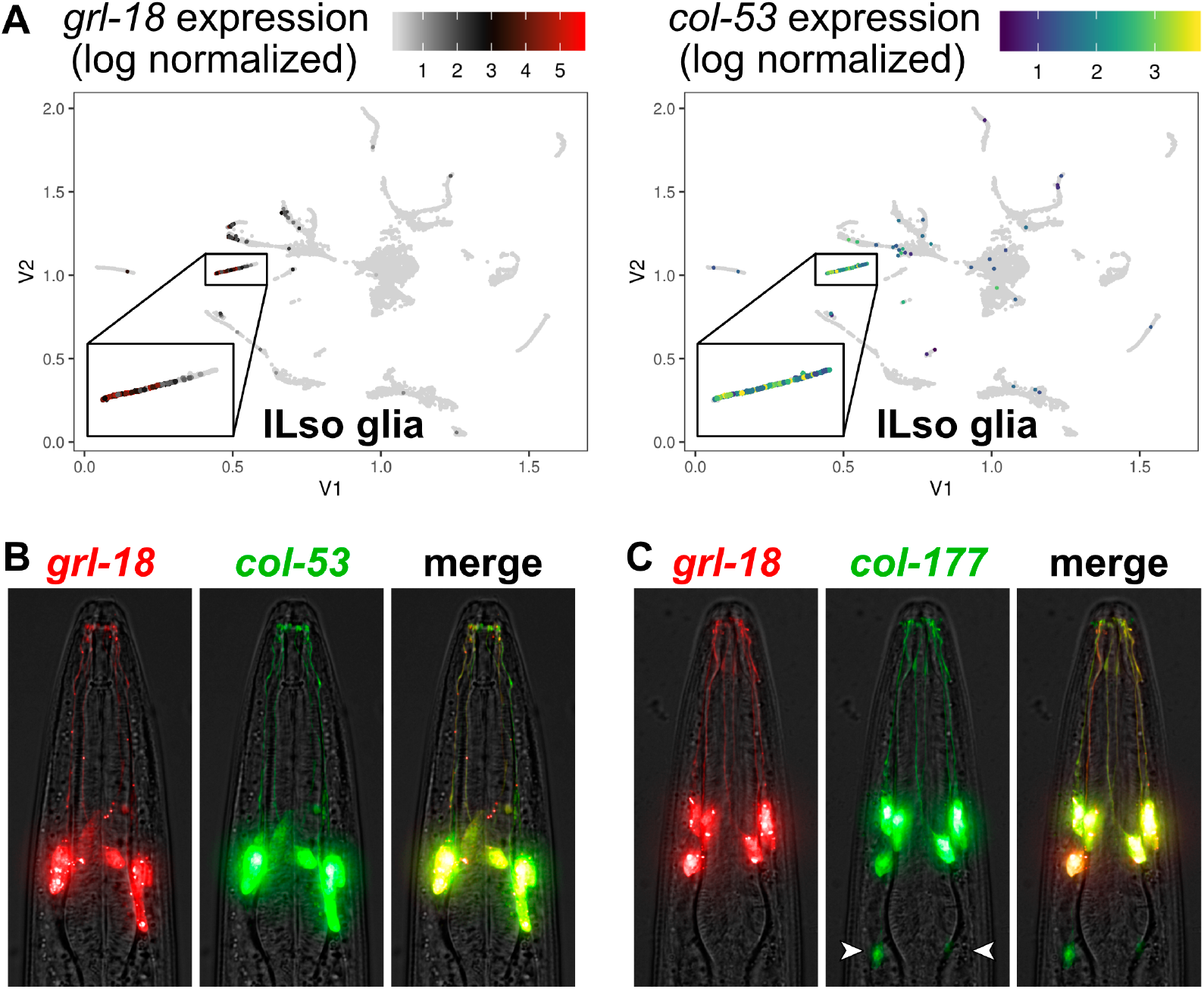
Prospective identification of novel cell-type-specific promoters for ILso glia. (A) Cell cluster plots illustrating the expression of *grl-18* and *col-53* in embryonic glia and excretory cells from Packer et al., 2019. Each point represents an individual cell. The color indicates the relative expression level of each gene. For *grl-18*, low expression is black and high expression is red. For *col-53*, low is blue and high is green/yellow. Gray indicates no detected transcripts for the gene of interest. Inset box, the cell cluster predicted to include ILso glia. Cell cluster plot for *col-177* is in Supp. Fig. S1A. (B, C) Merged brightfield and fluorescence images of an animal expressing *grl-18*pro:mApple together with (B) *col-53*pro:GFP or (C) *col-177*pro:GFP, showing that these markers are expressed in the same cells. *col-177* is also faintly expressed in cells with a glial morphology, tentatively identified as OLL socket glia in the head (arrowheads, see Supp. Fig. S1B) and ADE and PDE socket glia in the body.

We created *col-53* and *col-177* reporters consisting of 733 bp or 2.75 kb promoter regions upstream of their respective translation start sites fused to GFP. We generated animals with a *col-53*pro:GFP or *col-177*pro:GFP extrachromosomal array and an integrated *grl-18pro*:mApple transgene. We found that *col-53* and *col-177* are each expressed in the same six glial cells as *grl-18* (Fig. 5B, C), demonstrating that they are novel cell-type-specific promoters for ILso glia. For *col-177*, we found that it is also expressed in two additional cells in the head that sit posteriorly to the ILso glia and extend processes to the tip of the nose. Based on morphology, position, symmetry, and the association of OLL neuron sensory dendrites with their distal endings (Supp. Fig. S1B), we have tentatively identified these cells as the lateral OLL/Rso glia, for which there is currently no marker. We also observe faint expression in cells in the body whose morphology, position, and symmetry is consistent with the anterior and posterior deirid (ADE, PDE) socket glia.

Our findings provide proof-of-principle that single-cell transcript profiling enables prospective identification of novel cell-type-specific glial promoters. Indeed, we have also found that this approach can be used to identify promoters that are expressed in combinations of glia (*mam-5*, Supp. Fig. S2A, B) or that are likely to be expressed in other discrete glial types (AMso, Supp. Fig. S2C). Markers for the remaining unexplored glia will be essential for studies that systematically examine the roles of glia in the diverse sense organs of *C. elegans*. Furthermore, it will be extremely valuable to identify glial promoters that are expressed early in development or whose expression changes with environmental context, such as neuronal activity. Glial markers expressed in embryos will allow us to visualize and track the birth, migration, and morphogenesis of glia during nervous system assembly (Rapti et al., 2017). Meanwhile, glial markers whose expression is regulated by the environment will provide insight on how glia adapt and potentially alter interactions with their associated neurons to drive appropriate behaviors. These rich datasets and tools will open up new avenues for discovery in the field of glial biology.

## MATERIALS AND METHODS

### Strains and plasmids

All strains were grown at 20°C on nematode growth media (NGM) with *E. coli* OP50 bacteria (Brenner, 1974). Transgenic strains generated in this study were constructed in an N2 background. N2 animals were injected with 25-75 ng/μL per plasmid at a final concentration of 100 ng/μL DNA. Additional information on strains and plasmids used in this study are in Supplemental Table S1.

### Fluorescence microscopy and image processing

L2 to L3 animals were selected based on size and the extent of vulva development on the day of imaging. Animals were washed and immobilized in M9 solution containing 50 mM sodium azide and mounted on 2% agarose pads. Image stacks were collected on a DeltaVision Core imaging system (Applied Precision) with a UApo 40x/1.35 NA, PlanApo 60x/1.42 NA, or UPlanSApo 100x/1.40 NA oil immersion objective and a CoolSnap HQ2 camera. Images were subsequently deconvolved using Softworx (Applied Precision) and maximum intensity projections were generated in ImageJ. The brightness and contrast of each projection were linearly adjusted in Affinity Photo 1.7.3. Fluorescent signals were pseudo-colored, and merged images were generated using the Screen layer mode in Affinity Photo 1.7.3. When the whole animal is shown, a composite of multiple fields was manually stitched together. The maximum intensity projections of the stitched images were generated with the same optical stacks across the whole animal, except for *F16F9.3* and *grl-2*, for which different optical stacks were used to generate the maximum intensity projections of the head and tail, because the head and tail glia are in different planes. Additionally, for *grl-2*, the head was split into two regions and different optical stacks were used to generate the maximum intensity projections to resolve both the AMso glia and the excretory cells separately. All figures were assembled in Affinity Designer 1.7.3. For quantification of cell fluorescence expression, FIJI software (Schindelin et al., 2012) was used to measure the integrated density of GFP or RFP in the cell body and the average of three background regions in the head was subtracted.

### Assessment of glial marker consistency

To assess the consistency of each glial marker, L2 to L3 transgenic animals were washed and immobilized in M9 solution containing 50 mM sodium azide and mounted on 2% agarose pads. Fluorescently labeled glial cells were scored visually across optical stacks with a Deltavision Core imaging system (Applied Precision) and a PlanApo 60x/1.42 NA oil immersion objective.

### Analysis of single-cell RNA sequencing data

To identify novel markers for ILso glia, we used the “Differential Expression” feature on the VisCello data explorer (Packer et al., 2019). We generated a list of genes that are highly enriched in the predicted ILso glia cluster compared to all other cell types in the single-cell RNA sequencing dataset. This was achieved by first selecting the following: Global dataset (Choose Sample), Cell type/subtype (Meta Class), and ILso (Group 1). Group 2 was left blank. Next, downsample cells was performed with the default settings (0.05 FDR cutoff, 200 Max Cell in subset, and 1000 Max Cell in background). Finally, differential expression (DE) analysis was run to generate a table of differentially expressed genes for the ILso glia cluster. A list of genes that are enriched in other glial types, including AMso glia (Supp. Fig. S2C), can be generated by replacing ILso with the glial type of interest (Group 1).

The graphical plots of *grl-18* and *col-53* or *col-177* expression (Fig. 5A, Supp. Fig. S1A) were generated using the online VisCello data explorer (https://cello.shinyapps.io/celegans_explorer/). The “Cell Type” explorer was used to select the glia and excretory cell subset, and data was displayed using the “UMAP-2D [Paper]” projection colored by cell type/subtype. Separate searches for *grl-18* and *col-53* or *col-177* were used to generate a plot of the glia and excretory cell subset showing the relative expression levels of each gene using the black_red palette for *grl-18* and viridis palette for *col-53* or *col-177*. Inset boxes in Fig. 5A and Supp. Fig. S1A are magnifications of the predicted ILso cluster, which contains 163 cells.

## ACKNOWLEDGMENTS

We thank Shai Shaham and members of the Shaham laboratory, Meera Sundaram, and John Murray for sharing information and advice on glial-specific promoters and for providing strains and plasmids, and John Murray for help using the VisCello data explorer. We thank members of the Heiman laboratory for unpublished work further characterizing candidate promoters, including Ian McLachlan (*itx-1*), Elizabeth Cebul (*grl-18, delm-1, delm-2*), and Karolina Mizeracka (*grl-2, lin-48*). Some strains were provided by the CGC, which is funded by National Institutes of Health Office of Research Infrastructure Programs (P40 OD010440). This work was supported by NIH R01NS112343 to M.G.H.

