## Supplemental Data for "Cell-type-specific promoters for *C. elegans* glia"

### Supplemental Table S1. Strains and plasmids

#### (A) Strains from previous studies

| Strain | Genotype | Figures | References |
|---|---|---|---|
| OS1579 | *nsEx856* [*F16F9.3*pro:GFP, *rol-6(su1006)*] | 2A, 2B | Bacaj et al., 2008 |
| OS1585 | *nsEx864* [*F11C7.2*pro:GFP, *rol-6(su1006)*] | 2A; quantification described in Results | Bacaj et al., 2008 |
| VPR839 | *irIs67* [*hlh-17*pro:GFP, *unc-119(+)*]; *unc-119(ed4)* | 2A, 2B; quantification described in Results | Stout et al., 2013 |
| OS4260 | *nsIs198* [*mir-228*pro:GFP] | 3A, 3C | Rapti et al., 2017 |
| CHB1549 | *hmnIs13* [*pPD95.75F16F9.3*pro:mCherry, *grl-2*pro:YFP, *gcy-8*pro:myrCFP] | quantification described in Results | Mizeracka et al., 2019 |
| CHB3746 | *hmnEx2122* [*grl-2*pro:CFP, *grl-18*pro:YFP, *rol-6(su1006)*] | 2A, 2B | Mizeracka et al., 2019 |
| OS146 | *lin-15(n765) ; nsIs22* [*lin-15*(+), *ver-1*pro:GFP] | 4B, 4D | Popovici et al., 2002 |

#### (B) Strains from this study

| Strain | Genotype | Figures |
|---|---|---|
| CHB3829 | *hmnIs82* [*grl-18*pro:GFP] | quantification described in Results |
| CHB3936 | *hmnIs82* [*grl-18*pro:GFP]; *hmnEx176* [*klp-6*pro:mCherry] | 2A, 2C |
| CHB1641 | *hmnEx877* [*delm-1pro*:GFP, *itx-1*pro:mApple, *rol-6(su1006)*] | 3A, 3B |
| CHB3981 | *hmnEx2212* [*itx-1*(2977 bp)pro:GFP, *rol-6(su1006)*] | 3A, 3B |
| CHB921 | *nsIs53* [*vap-1*pro:RFP] | 4A, 4C |
| CHB4037 | *hmnIs47* [*grl-18*pro:mApple]; *hmnEx2229* [*col-53pro*:GFP; *rol-6(su1006)*] | 5B |
| CHB3842 | *hmnIs47* [*grl-18*pro:mApple]; *hmnEx2171* [*col-177*pro:GFP, *rol-6(su1006)*] | 5C |
| CHB3986 | *hmnEx2215* [*col-177*pro:GFP; *ser-2prom3*:mCherry] | S1B |
| CHB4038 | *hmnIs47* [*grl-18*pro:mApple]; *hmnEx2207* [*mam-5*pro:GFP; *rol-6(su1006)*] | S2B |

#### (C) Plasmids from this study

| Plasmid | Description | Notes |
|---|---|---|
| pEL141 | *grl-18*pro:mApple | uses a 2962 bp genomic DNA fragment upstream of the translation start site of *grl-18* (gift of Elizabeth Cebul) |
| pWF2 | *col-177*pro:GFP | uses a 2750 bp genomic DNA fragment upstream of the translation start site of *col-177* |
| pWF3 | *mam-5*pro:GFP | uses a 620 bp genomic DNA fragment upstream of the translation start site of *mam-5* |
| pWF25 | *col-53*pro:GFP | uses a 733 bp genomic DNA fragment upstream of the translation start site of *col-53* |

**Supplemental Table S2**

| Gene | Glial type (number of cells) | Glial cells expressing transgene ±SD | *N* |
|---|---|---|---|
| *F16F9.3* | AMsh (2) | 1.96 ± 0.2 | 25 |
| | PHsh (2) | 2.0 ± 0.0 | |
| *F11C7.2* | AMsh (2) | 1.70 ± 0.5 | 23 |
| *hlh-17* | CEPsh (4) | 4.0 ± 0.0 | 22 |
| *grl-2* | AMso (2) | 2.0 ± 0.0 | 23 |
| | PHso1 (2) | 2.0 ± 0.0 | |
| | PHso2 (2) | 2.0 ± 0.0 | |
| *grl-18* | ILso (6) | 6.0 ± 0.0 | 28 |
| *delm-1* | Multiple types | 6.4 ± 1.2 | 20 |
| *itx-1* | Multiple types | 9.4 ± 2.3 | 24 |

**Supp. Table S2. Consistency of recommended glial markers**

All transgenes were genomically integrated except *F11C7.2*, *delm-1*, and *itx-1* for which an extrachromosomal array was used, which may contribute to their less consistent expression.

**Supplemental Figure S1**

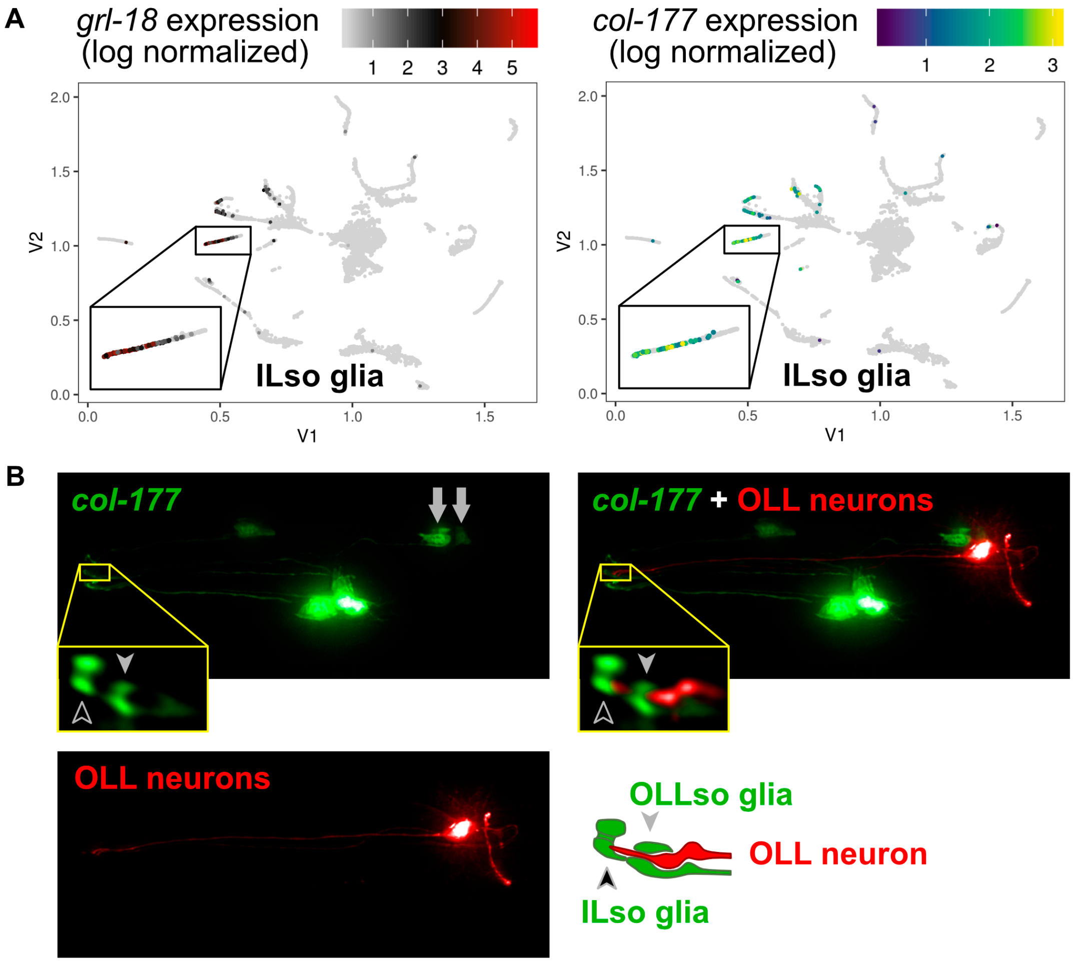


**Supp. Fig. S1. Expression pattern of *col-177***

(A) Cell cluster plot illustrating the expression of *grl-18* and *col-177* in embryonic glia and excretory cells from Packer et al., 2019. Each point represents an individual cell. The color indicates the relative expression level of each gene. For *grl-18*, low expression is black and high expression is red. For *col-177*, low is blue and high is green/yellow. Gray indicates no detected transcripts for the gene of interest. Inset, cell cluster predicted to include ILso glia. (B) *col-177*pro:GFP is expressed brightly in ILso (see Fig. 5C) and faintly in two cells with glial morphology on either side of the head (arrows) that appear to wrap the OLL dendritic endings and thus may be OLLso glia. *col-177*pro:GFP, green; *ser-2prom3*:mCherry in OLL neurons, red. ILso glial pore, black arrowhead; OLLso glial pore, gray arrowhead.

**Supplementary Figure S2**

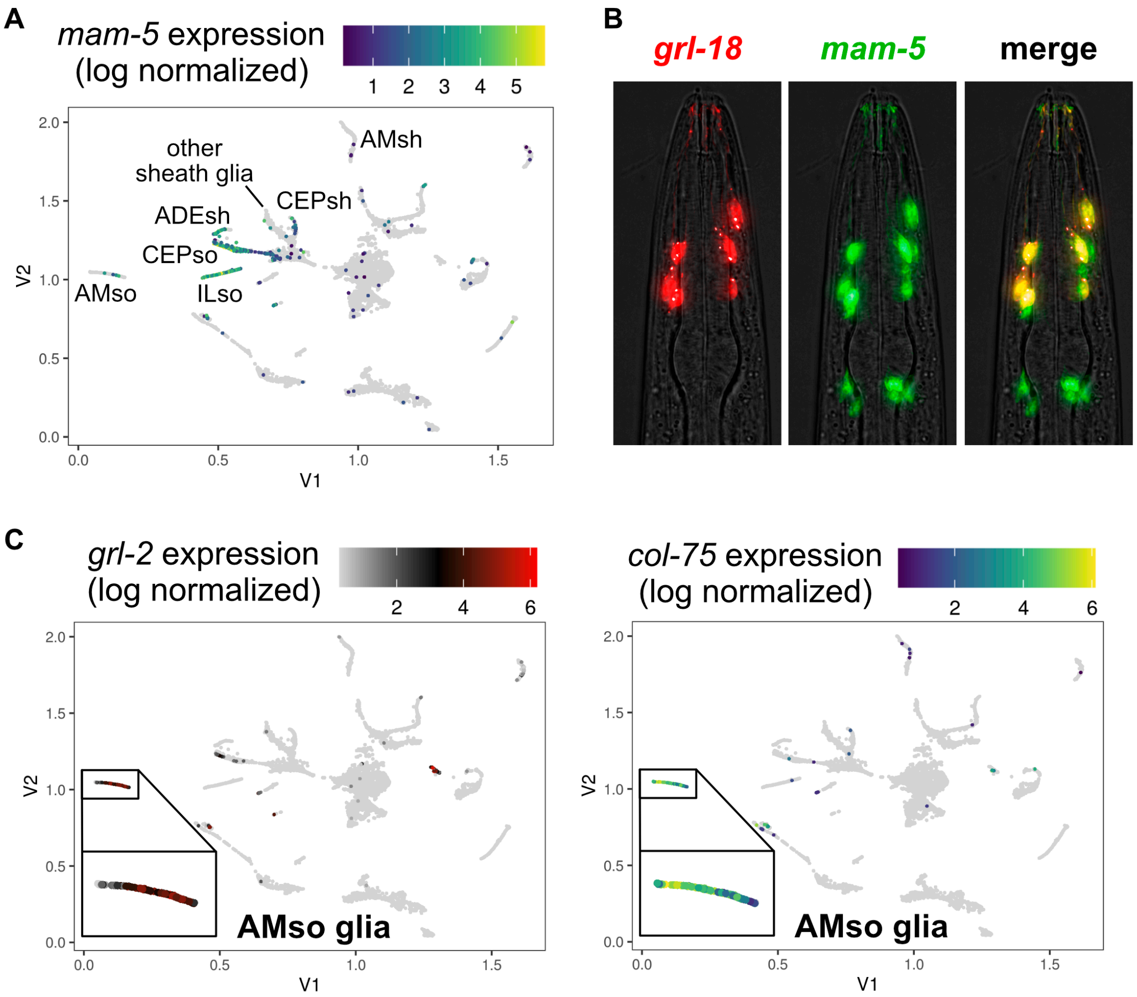


**Supp. Fig. S2. Prospective identification of promoters for other glial types**

(A) Cell cluster plot illustrating the expression of *mam-5* in embryonic glia and excretory cells from Packer et al., 2019, as in Supp. Fig. S1. Clusters previously nominated to correspond to specific glial types are indicated. CEPso cluster may contain other socket glia. *mam-5* is predicted to be expressed in ILso glia and other glial types. (B) *mam-5*pro:GFP (green) is expressed with *grl-18*pro:mApple (red) in ILso glia and other head cells with glial morphology. (C) A cell cluster previously identified as AMso glia is enriched for the known marker *grl-2* (left) as well as several collagen genes including *col-75* (right), *col-174* and *col-187*, demonstrating how this approach can be used to identify novel candidate markers of other glial types.
